# Neural coding of leg proprioception in *Drosophila*

**DOI:** 10.1101/274498

**Authors:** Akira Mamiya, Pralaksha Gurung, John Tuthill

## Abstract

Animals rely on an internal sense of body position and movement to effectively control motor behavior. This sense of proprioception relies on diverse populations of mechanosensory neurons distributed throughout the body. However, little is known about how proprioceptor neurons collectively encode sensory stimuli. Here, we investigate neural coding of leg proprioception in *Drosophila*, using *in vivo* two-photon calcium imaging of proprioceptors during controlled movements of the fly tibia. We found that the axons of leg proprioceptors are organized into distinct functional projections that contain topographic representations of specific kinematic features. Using subtype-specific genetic driver lines, we show that one group of axons encodes tibia position (flexion/extension), another encodes movement direction, and a third encodes bidirectional movement and vibration frequency. Thus, proprioceptive sensing of a single leg joint is mediated by multiple subtypes of specialized sensory neurons. This architecture may help to maximize information transmission, processing speed, and robustness, which are critical for feedback control of the limbs during locomotion.

## Introduction

Proprioception, the internal sense of body position and movement (Sherrington, 1906), is essential for the neural control of motor behavior. Sensory feedback from proprioceptive sensory neurons (i.e., proprioceptors) contributes to a wide range of behaviors, from regulation of body posture (Hasan and Stuart, 1988; Zill et al., 2004), to locomotor adaptation (Bidaye et al., 2018; Lam and Pearson, 2002) and motor learning (Isakov et al., 2016; Takeoka et al., 2014). But despite the fundamental importance of proprioception to our daily experience, it is perhaps the most poorly understood of the primary senses.

Proprioceptors are found in nearly all motile animals, from pubic lice (Graber, 1882) to sperm whales (Sierra et al., 2015). Single-unit electrophysiological recordings have revealed that proprioceptors can vary widely in their mechanical sensitivity and stimulus tuning, even within a single limb segment or muscle. For example, most vertebrate muscles contain two distinct classes of proprioceptive organs that are innervated by specialized sensory neurons: muscle spindles detect muscle length and contraction velocity, while golgi tendon organs detect mechanical load (Proske and Gandevia, 2012; Windhorst, 2007). Insects also possess proprioceptor subtypes that encode joint position, velocity, and load (Burrows, 1996; Tuthill and Wilson, 2016), suggesting that the nervous systems of invertebrates and vertebrates may have arrived at similar evolutionary solutions to a set of common sensorimotor constraints (Tuthill and Azim, 2018).

Although individual proprioceptors may be narrowly tuned, most body movements are likely to drive activity in large numbers of proprioceptors (Proske and Gandevia, 2012; Zill et al., 2004). Characterizing the population-level structure of proprioceptive encoding has been challenging, due to the technical difficulty of recording from multiple neurons simultaneously, or identifying the same neurons across individuals. It has also been difficult to identify spatial structure or topography within proprioceptive populations from recordings of single neurons. However, a detailed understanding of proprioceptive population coding is important for understanding the function of downstream circuits, and identifying the role of proprioceptive feedback in the neural control of movement.

Here, we combine genetic tools, 2-photon calcium imaging, and novel experimental setup to investigate the anatomy and function of a proprioceptor population from the leg of the fruit fly, *Drosophila*. Embedded within the femur of the insect leg is a cluster of proprioceptor cell bodies known collectively as the femoral chordotonal organ (FeCO; Field and Matheson, 1998). Experimental manipulation of the FeCO in locusts and stick insects has revealed a critical role for these proprioceptors during behaviors that require precise control of leg position, such as walking (Bässler, 1988) and targeted reaching (Page and Matheson, 2009). Single-unit electrophysiological recordings in these larger insect species have found that FeCO neurons monitor the position and movement of the femur-tibia joint, and that FeCO neurons may be narrowly tuned to specific kinematic features (Field and Pflüger, 1989; Hofmann et al., 1985; Kondoh et al., 1995; Matheson, 1990; Matheson, 1992; Stein and Sauer, 1999; Zill, 1985). However, it has been challenging to integrate data from single neurons to understand the topographic structure and function of the FeCO population. In *Drosophila*, where genetic tools enable systematic dissection of neuronal populations, the anatomy and physiology of FeCO neurons have not previously been investigated.

Using *in vivo* population-level calcium imaging of FeCO axons during controlled leg manipulations, we first mapped the spatial organization of proprioceptive signals in the fly ventral nerve cord (VNC). We then identified genetically and anatomically distinct FeCO subtypes that encode specific kinematic features, including position, movement direction, and movement frequency. Overall, our results reveal the basic architecture and neural code for a key leg proprioceptive system in *Drosophila*. These findings provide a framework for understanding how specialized proprioceptive feedback signals enable robust and accurate motor control.

## Results

### Methods to record and map proprioceptive signals in Drosophila

Positioned in the proximal femur of each *Drosophila* leg is a femoral chordotonal organ (FeCO) that contains ∼135 cell bodies (**Figure 1A**). The dendrites of the FeCO neurons are connected to the cuticle and surrounding muscles by attachment cells and tendons (Shanbhag et al., 1992), and the axons of FeCO neurons project through the leg nerve into the ventral nerve cord (VNC; Phillis et al., 1996; Smith and Shepherd, 1996). Most FeCO axons arborize within the VNC neuropil (**Figure 1A, D**), with only 3-4 cells from each leg projecting directly to the brain (Tsubouchi et al., 2017).

**Figure 1.**
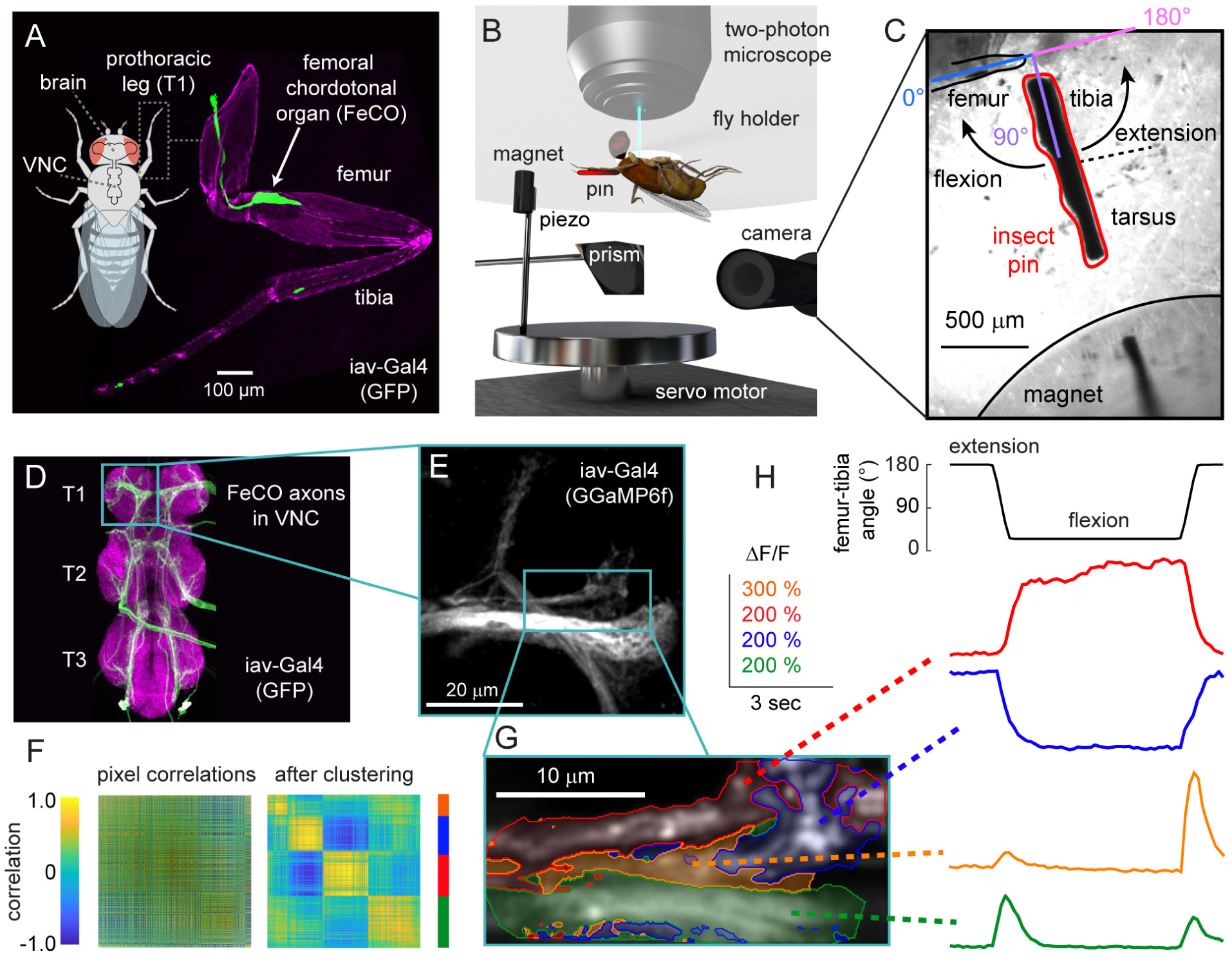
Investigating proprioceptive tuning of the *Drosophila* femoral chordotonal organ (FeCO). **A.** A confocal image of the front (T1) leg of *Drosophila melanogaster*, showing the location of the FeCO cell bodies and dendrites (green). Magenta is auto-fluorescence from the cuticle. **B.** An experimental setup for 2-photon calcium imaging from the axons of FeCO neurons while controlling and tracking the femur-tibia joint. To control joint angle, we glued a pin to the tibia and positioned it using a magnet mounted on a servo motor. We vibrated the tibia with a piezoelectric crystal fixed to the magnet. We backlit the tibia with an IR LED and recorded the tibia position from below using a prism and high-speed video camera. **C.** An example frame from a video used to track joint angle. The pin is painted black to enhance the contrast against the background. **D.** FeCO axon terminals (green) in the fly ventral nerve cord (VNC). Magenta is a neuropil stain (nc82). Teal box indicates region imaged by the 2-photon microscope. **E.** 2-photon image of FeCO axon terminals expressing GCaMP6f driven by *iav-Gal4*. The teal box indicates the region imaged in the example recording shown in panels G-H. **F.** A cross-correlation matrix showing pixel-to-pixel correlations of the changes in GCaMP6f fluorescence (ΔF/F) during an example trial shown in panels G-H. Left: the cross-correlation matrix before correlation based clustering. Right: the matrix after the clustering. The colors on the right of the correlation matrix correspond to the cluster colors used in G-H. **G.** Image of GCaMP6f fluorescence showing the recording region for the example trial shown in F and H. Each group of pixels is shaded according to its cluster identity. The colors correspond to the ΔF/F traces shown in H. **H.** Calcium signals from different clusters of pixels during an example trial. Each trace shows the changes in GCaMP6f fluorescence (ΔF/F) for the groups of pixels shaded with the same color in the panel G.

To investigate proprioceptive signal encoding of the FeCO population, we recorded the activity of proprioceptor axons while manipulating the joint angle between the femur and tibia of the fly’s right front (T1) leg (**Figure 1B, C**). We used the Gal4-UAS system (Brand and Perrimon, 1993) to express a genetically encoded calcium indicator, GCaMP6f (Chen et al., 2013) in the majority of FeCO neurons (*iav-Gal4*; Kwon et al., 2010) and recorded their calcium activity *in vivo* with two-photon calcium imaging (**Figure 1B**). For fast and accurate positioning of the leg, we designed a magnetic control system that allowed us to manipulate a pin glued to the fly’s tibia using a servo-actuated magnet (**Figure 1B**). The femur and proximal leg joints were fixed to the fly holder with UV-cured glue. We continuously recorded the position of the tibia using an IR-sensitive video camera at 180 - 200 Hz (**Figure 1C**), and automatically tracked its orientation to calculate the femur-tibia joint angle.

To identify proprioceptor projections with shared functional tuning, we recorded calcium activity of FeCO axons as we swung the tibia from flexion to extension (**Figure 1F-H**). To categorize calcium activity in an unbiased manner, we then calculated pairwise correlations between the calcium signal (ΔF/F) in each pixel, and performed k-means clustering on the resulting correlation matrix. **Figure 1F** shows an example of a pixel correlation matrix before and after clustering. In this trial, we identified four groups of pixels whose activity was highly correlated. Although this correlation-based clustering does not impose any spatial restrictions, we found that the pixels that clustered together based on their activity were also grouped together spatially (**Figure 1G-H**). This example illustrates that clustering of pixel correlations is sufficient to identify groups of FeCO axons that encode distinct proprioceptive stimulus features.

### Functional organization of FeCO axons in the fly ventral nerve cord (VNC)

To identify functional subtypes within the FeCO population, we recorded calcium signals from four regions of interest that encompassed the different axon projections in the VNC (**Figure 2**). For these experiments, we first clustered similarly responding pixels within each trial to identify the different response types. Then, we combined the responses recorded from the same region in different flies and again grouped them using correlation based k-means clustering (**Figure 2A-D: middle columns**).

**Figure 2.**
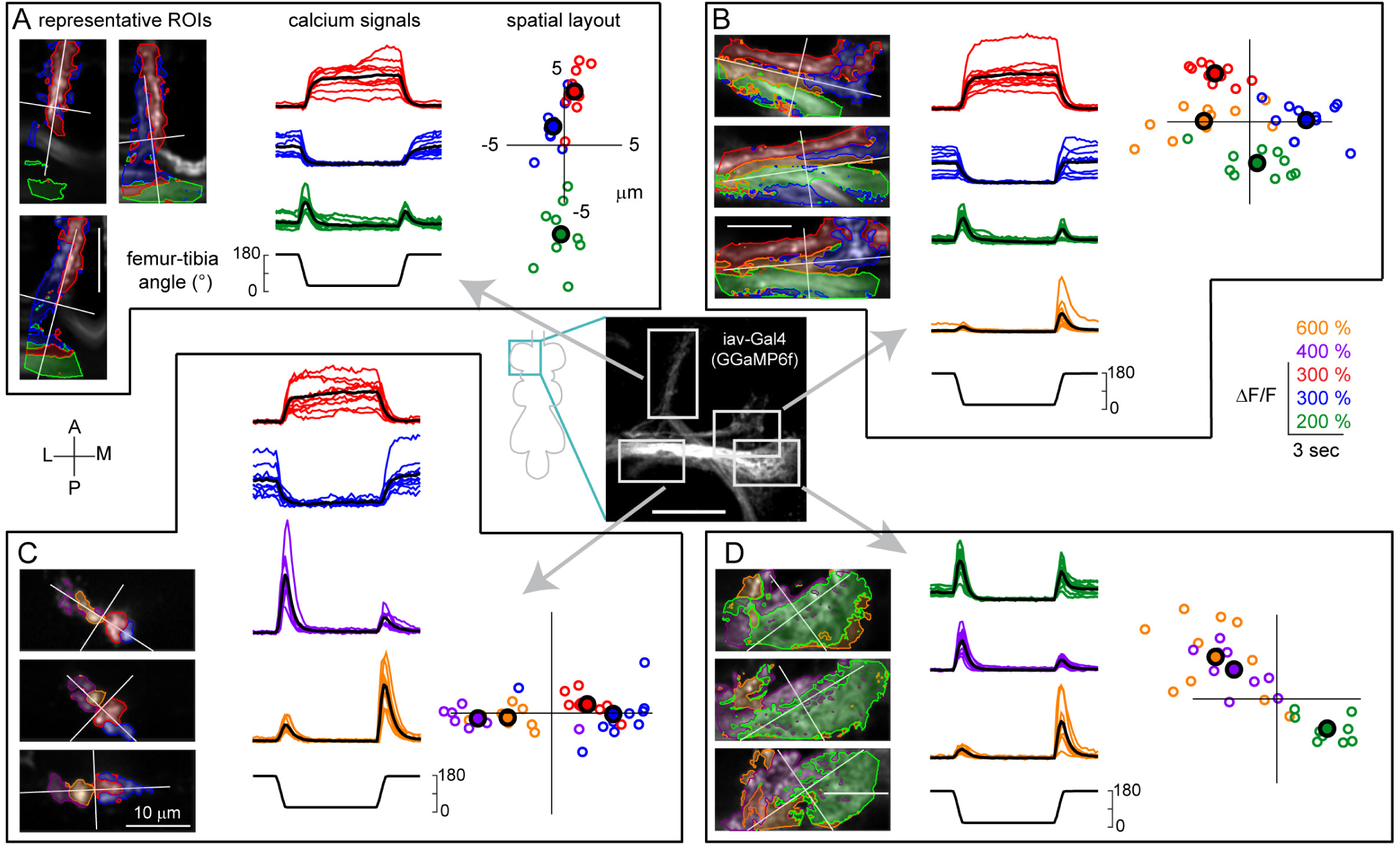
FeCO axons encode distinct proprioceptive features. Each panel (A-D) shows calcium signals recorded from FeCO axons (*iav-Gal4*; *UAS-GCamp6f*) within a different region of interest. **A**. Imaging from an anterolateral region. *Left column*: example images of GCaMP6f fluorescence from three flies. In each fly, we categorized the pixels into three clusters (as in Figure 1F). The red and blue regions are tonically active when the tibia is flexed (red) or extended (blue). The green region responds phasically during tibia movement. The color scheme is maintained throughout the figure. A white cross represents the center of the recording region, aligned to the long axis of each axon projection. *Middle column*: Calcium signals for each group of pixels recorded from multiple flies. Red, blue, and green lines show GCaMP6f fluorescence (ΔF/F) during the trial for each cluster of pixels (red: n= 10 flies, blue: n = 10, green: n = 9). Black lines show the response average. *Right column*: the location of each cluster relative to the center of the recording region across flies. The larger outlined circles represent the mean location of the center for each cluster. Each image was rotated to a common axis (see Methods for details). **B**. Imaging from an anteromedial region (red: n= 11 flies, blue: n = 11, green: n = 10, orange: n = 10). In addition to the three types of responses we identified in A, this region also contained a group of pixels that responded phasically during tibia extension (orange). **C**. Imaging from a posterolateral region (red: n= 10 flies, blue: n = 10, purple: n = 8, orange: n = 9). Here, we identified pixels that responded phasically during joint flexion (purple). **D**. Imaging from a medial region (green: n= 10 flies, purple: n = 11, orange: n = 10). The scale bar in the center image is 20 μm.

We identified five basic types of responses within the FeCO axons: two tonic (non-adapting) and three phasic (adapting). Pixels that responded tonically increased their activity when the tibia was either flexed (blue) or extended (red). Phasic pixels increased their activity transiently during the movement phase of the swing stimulus; one group responded to movements in both directions (green), while the two other groups responded in a directionally selective manner to either flexion (orange) or extension (purple). Each functional response type was consistently located in similar positions across multiple flies (**Figure 2A-D: right columns; Figure S1D-G**). This analysis suggests that FeCO neurons that encode distinct kinematic features (i.e., tibia position, movement, and direction) are grouped into functional projections within the VNC.

### Specific genetic driver lines delineate club, claw, and hook proprioceptors

Population-level imaging from FeCO neurons with a broad driver line (*iav-Gal4*) revealed that different axon projections have distinct proprioceptive tuning and are systemically organized across flies (**Figure 2**). We next sought to identify more specific Gal4 driver lines that would provide a genetic handle for these functional subtypes and enable more fine-grained analysis of proprioceptor anatomy and function. We screened through Gal4 lines in the Janelia FlyLight collection (Jenett et al., 2012) for lines that drove expression in each axon projection. From this screen, we identified three Gal4 lines that labeled the cardinal FeCO axon projections (**Figure 3**).

**Figure 3.**
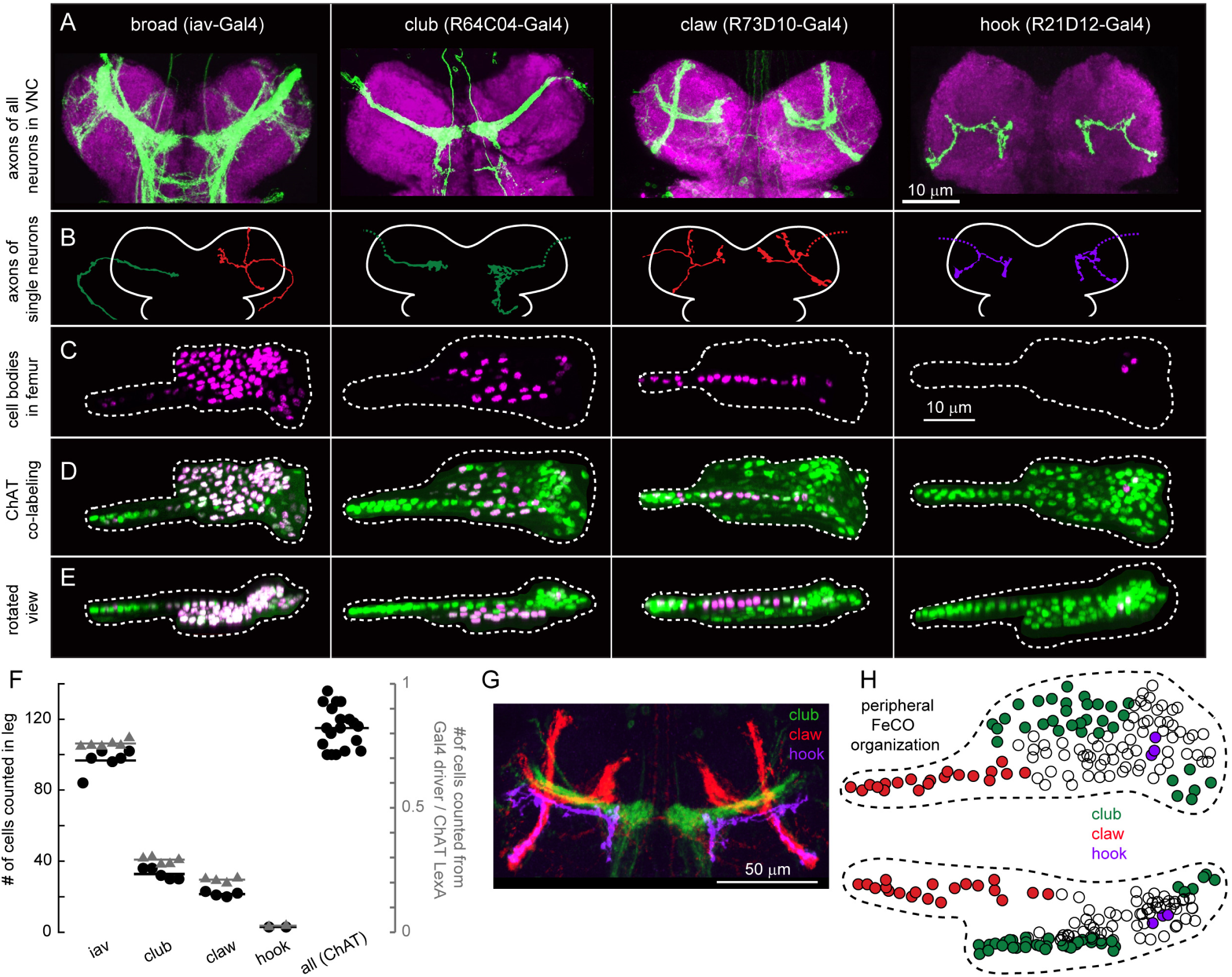
Organization of genetically-defined FeCO neuron subtypes in the VNC and leg. **A**. Four Gal4 lines label subsets of FeCO axons in the VNC. Green: GFP driven by each Gal4, magenta: nc82 neuropil staining. Scale bar is 50 μm. **B**. Example morphologies of single FeCO neurons driven by each Gal4 line, traced from images obtained by stochastic labeling with the multi-color flp-out technique. Dotted lines indicate severed axons. **C**. FeCO cell bodies are clustered in characteristic locations in the fly leg. Cell bodies were labeled with *UAS-Redstinger* driven by each Gal4 line. Scale bars are 10 μm. **D**. Co-labeling of FeCO cell bodies from each Gal4 line (as in C), with a green reference that labels all FeCO neurons (*Chat-Lexa*; *UAS-nlsGFP*). **E**. Same as D, but viewed from the dorsal side. **F**. Number of neurons labeled by each Gal4 line shown in A (black circles) and the ratio of neurons labeled by each Gal4 line compared to those labeled by *ChAT-LexA* in the same leg (blue triangles); *iav*, n = 6 flies, club, n = 5 flies, claw, n = 4 flies, hook, n = 2 flies, *ChaT*, n = 19. **G**. *in silico* overlay of the axon projections of the club, claw, and hook neurons in the VNC. **H**. A schematic of the FeCO in the femur, showing the location of cell bodies labeled by each Gal4 line.

The first driver line (*R64C04-Gal4*) labels axons that run laterally through the center of the leg neuropil and form an arborization in the shape of a *club* (**Figure 3A, 2^nd^ column;** nomenclature from Phillis et al., 1996). The majority of the club axons terminate near the midline, although some also project toward the brain or other VNC segments. The second line (*R73D10-Gal4*) labels a group of axons shaped like a *claw* (Phillis et al., 1996). Upon entering the VNC from the leg nerve, this projection splits into three smaller branches (**Figure 3A, 3^rd^ column**). We refer to these as the X, Y, and Z branches: the X branch of the claw continues to run parallel to the club, while the Y branch projects anteriorly, and the Z branch projects dorsally. The third line (*R21D12-Gal4*) labels axons that project along the Z branch of the claw, with only a small protuberance along the Y branch, and extend a longer arborization toward the midline, just anterior to the club (**Figure 3A, 4^th^ column**). We call this projection the *hook* due to its resemblance to a lumberjack’s peavey hook.

To characterize the anatomy of single neurons within each driver line, we used the multicolor flip-out technique (Nern et al., 2015) to stochastically label single cells, and manually traced their morphology (**Figure 3B**). Every neuron we traced from the claw Gal4 line (n=3) had a similar morphology, including projections to all three branches (X, Y, and Z) of the claw. Similarly, all the cells we traced from the hook Gal4 line (n=2) were morphologically similar, with each cell projecting an axon along all branches of the hook. In contrast, we identified three unique morphologies within the club Gal4 line (n=14), indicating that there may exist further functional subdivisions within the club.

For each Gal4 line, we investigated the number and distribution of proprioceptor cell bodies in the femur using confocal microscopy (**Figure 3C-E**). We drove expression of a nuclear-localized red fluorescent protein (UAS-RedStinger) with each Gal4 line, while expressing a nuclear-localized GFP in all FeCO neurons using ChAT-LexA line, which labels all leg mechanosensory neurons. (We were able to unambiguously identify FeCO neurons in ChAT-LexA based on their characteristic location in the leg.) In addition to the subtype-specific Gal4 lines, we performed similar co-labeling experiments using the broadly expressing line *iav-gal4*. Although *iav-Gal4* was previously thought to drive expression in all FeCO neurons, we found that it labeled ∼80% of the total population (∼136 neurons; **Figure 3F**).

The claw Gal4 line labeled ∼20 cell bodies (**Figure 3F**), which were located in a characteristic blade-shaped strip that extended along the long axis of the femur (**Figure 3C-E**). The club Gal4 line drove expression in ∼30 neurons (**Figure 3F**); these cell bodies were located in the proximal part of the FeCO (**Figure 3C-E**). Finally, the hook Gal4 line drove expression in only 3 FeCO neurons (**Figure 3F**), which were located along the FeCO’s ventral edge (**Figure 3C-E**). The cell body location for each subtype was consistent across flies. These results suggest that the cell bodies of FeCO neurons with distinct central projections are grouped in specific locations in the FeCO (**Figure 3H**). Although these three Gal4 lines label less than half of the total FeCO neuron population, computational registration to a standard VNC confirmed that these three subtypes span the cardinal FeCO projections labeled by *iav-Gal4* (**Figure 3G**).

### Calcium imaging from specific Gal4 lines shows that position, movement, and direction are encoded by separate proprioceptor subtypes

We next performed calcium imaging from the claw, club, and hook neurons. Claw neurons increased their activity tonically in response to either flexion or extension of the tibia (**Figure 4A**). Each branch of the claw axon projection had two sub-branches that responded either to flexion or extension, which we identified with the clustering methods described above (**Figure 4A**). The time courses of these responses were similar across the X, Y, and Z branches of the claw (**Figure 4A, Figure S2A**), consistent with our anatomical observation that each claw neuron innervates all three branches (**Figure 3B**). The spatial organization and functional tuning of claw neurons are also consistent with the position-tuned tonic responses recorded from population imaging (**Figure 2**).

**Figure 4.**
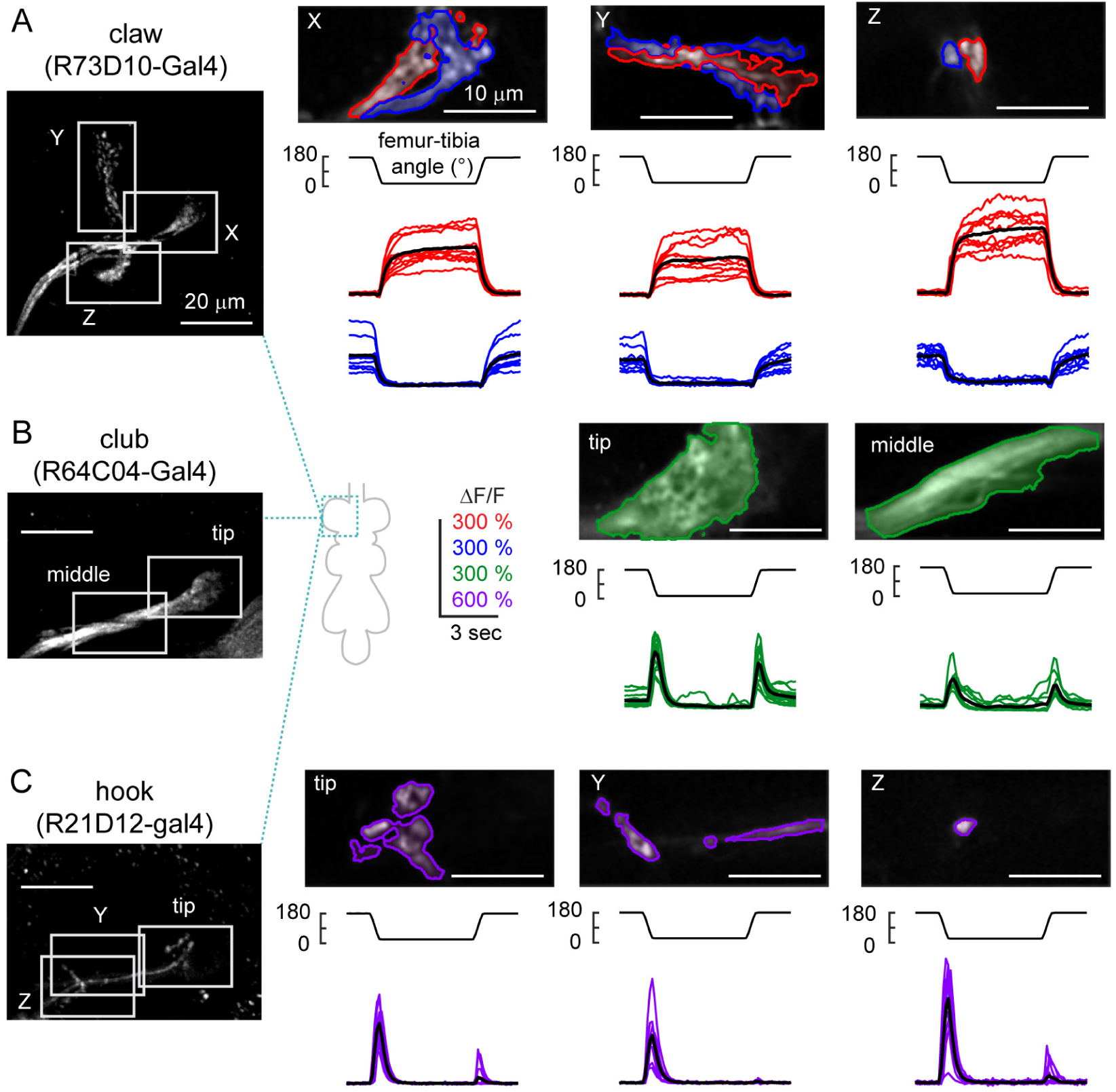
Three Gal4 lines delineate FeCO functional subtypes. **A.** Claw neurons encode tibia position, with distinct pixels responding to either flexion (red) or extension (blue). *Left*: The claw axon projection in the VNC visualized with GCaMP6f fluorescence driven by *R73D10-Gal4*. White rectangles represent the imaging locations for the X, Y, and Z branches shown in the right three columns. In each column, the top image shows a representative region of interest, with flexion-encoding pixels shaded in red, extension encoding pixels in blue. The Y branch is rotated 90 degrees clockwise. The bottom rows show changes in GCaMP6f fluorescence relative to the baseline (ΔF/F) recorded from each sub-region in different flies (n = 10 flies for each region). Thick black lines represent average responses. **B.** Same as A, but for movement-sensitive club neurons (*R64C04-Gal4*). The club neurons respond phasically to both the flexion and the extension of the joint (n = 14 flies for tip, 11 flies for middle) **C.** Same as A and B, but for directionally-tuned hook neurons (*R21D12-Gal4*). The hook neurons respond phasically to flexion of the joint, but not extension (n = 14 flies for tip and Z, 9 flies for Y).

Calcium signals in club axons increased phasically during both flexion and the extension of the tibia (**Figure 4B**). The amplitude and time course of these calcium signals were similar during extension and flexion, and across different regions of the club (**Figure 4B, Figure S2B**). These results are consistent with population imaging data (**Figure 2**), although the bidirectional response symmetry in **Figure 4B** suggests that the club clusters identified from population imaging (**Figure 2**) may have been partly contaminated by nearby axons from directionally selective neurons.

The axons of hook neurons increased their activity phasically during tibia flexion, but responded only weakly during extension (**Figure 4C**). We recorded from three different locations along the hook axon projection and found that the responses were similar in all three locations (**Figures 4C, Figure S2C**). The time course of these responses were similar to those of the flexion-tuned phasic responses we observed in population-imaging experiments (**Figure 2**).

Together, our experiments imaging from claw, club, and hook-specific Gal4 drivers reveal that these lines label distinct proprioceptor subtypes that encode different stimulus features. The tuning of these lines matches the functional and spatial organization we observed in population imaging experiments with a broad driver line (**Figure 2**), indicating that they represent major FeCO subtypes. The only response type that we failed to identify with a specific Gal4 line was the extension-tuned complement of the hook neurons (orange traces in **Figure 2**). Based on the location of extension-tuned pixels in population imaging experiments, we expect these neurons to have similar morphology and projections to the flexion-tuned hook neurons. Next, we will use a broader range of proprioceptive stimuli to characterize the encoding properties of each proprioceptor subtype in more detail.

### Claw neurons encode tibia position, club neurons encode bidirectional tibia movement, and hook neurons encode directional tibia movement

We used ramp-and-hold stimuli to more comprehensively explore the position-dependent tuning of each proprioceptor subtype (**Figure 5**). The tibia started at either a flexed (∼18°) or extended (∼180°) position, then moved in 18° steps to the opposite position. Claw neurons responded to these stimuli with tonic increases in calcium during either flexion (red) or extension (blue) of the tibia (**Figure 5A**). We identified flexion and extension-tuned pixels using the same clustering methods that we used for population imaging (**Figures 1, 2**). The responses within these two regions were highly position-dependent: the activity of extension-tuned pixels increased when the tibia was between 90° and 180°, while the activity of flexion-tuned pixels increased when the tibia was between ∼90° and 18°. Interestingly, neither group was active in the middle of the joint range, close to 90°. Overall, the activity of claw neurons increased as a relatively linear function of the femur-tibia joint angle, although we also observed some directional hysteresis (see below).

**Figure 5.**
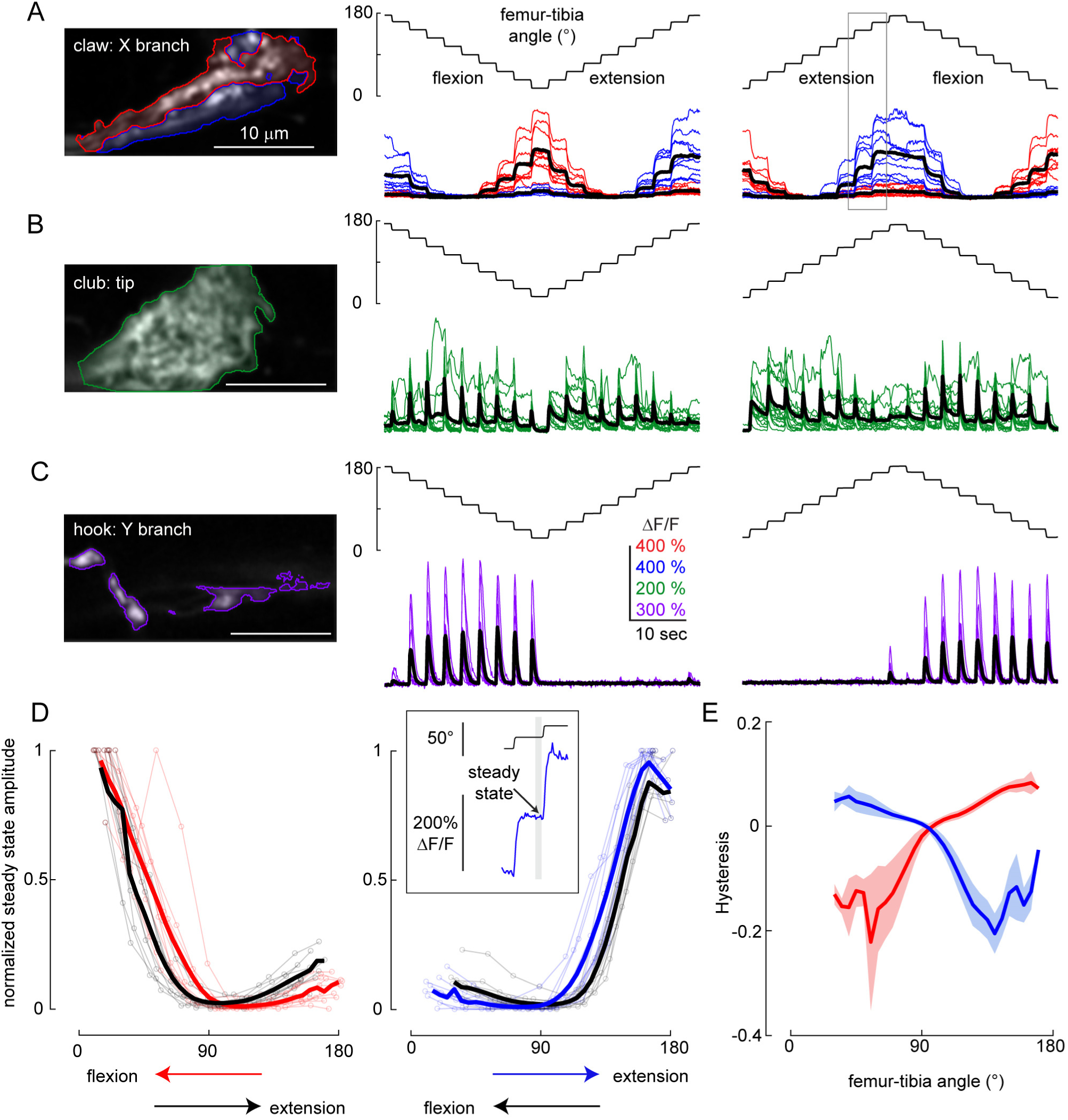
Claw neurons encode joint position, club neurons encode bidirectional movements, and hook neurons encode movement direction. **A.** Responses of position-encoding claw neurons (*R73D10-Gal4*) to ramp-and-hold stimuli. *Left column*: Average GCaMP6f fluorescence from the X branch of the claw projection where the example recordings were made (red: flexion encoding, blue: extension encoding). *Middle column*: Responses from the two regions to a ramp-and-hold stimulus that starts with the joint extended (n = 10 flies). *Right column*: Same as above, but starting with the joint flexed. The grey rectangle indicates the location of the trace shown in the top inset in D. **B**. Same as A, but for club neurons (*R64C04-Gal4*), which increase their activity phasically in response to each ramp step (n = 14 flies). **C**. Same as A and B, but for hook neurons (driven by *R21D12-Gal4*), which only respond during flexion (n = 9 flies). **D**. Calcium signals of claw neurons depend on movement history. *Left column:* steady state ΔF/F at different joint angles for the flexion activated (red: during flexion, black: during extension, thick lines: average response) sub-branches of the claw, normalized by the maximum peak response recorded in each fly (n= 10 flies). Steady-state responses were measured at the end of the hold step (top inset at right). In these recordings, flexion preceded extension. *Right column:* Same as the left column but for the extension activated (blue: during extension, black: during flexion, thick lines: average response) sub-branches of the claw (n = 10 flies). In these recordings, extension preceded flexion. **E**. Hysteresis (difference between the response to the activating direction and the non-activating direction) of the steady-state response for flexion (red) and extension (blue) activated sub-branches of the claw (thick lines: average response, shading: standard error of the mean).

Club neurons responded phasically to each tibia movement, regardless of movement direction (**Figure 5B**). Movement responses occurred across the joint angle range, but were slightly larger around 90° and smaller at full extension (180°). In addition to these bidirectional responses to movement steps, we occasionally observed tonic responses during the hold period, possibly due to low amplitude vibrations of the tibia (see **Figure 6**, below). These results confirm that club neurons respond to bidirectional tibia movement.

**Figure 6.**
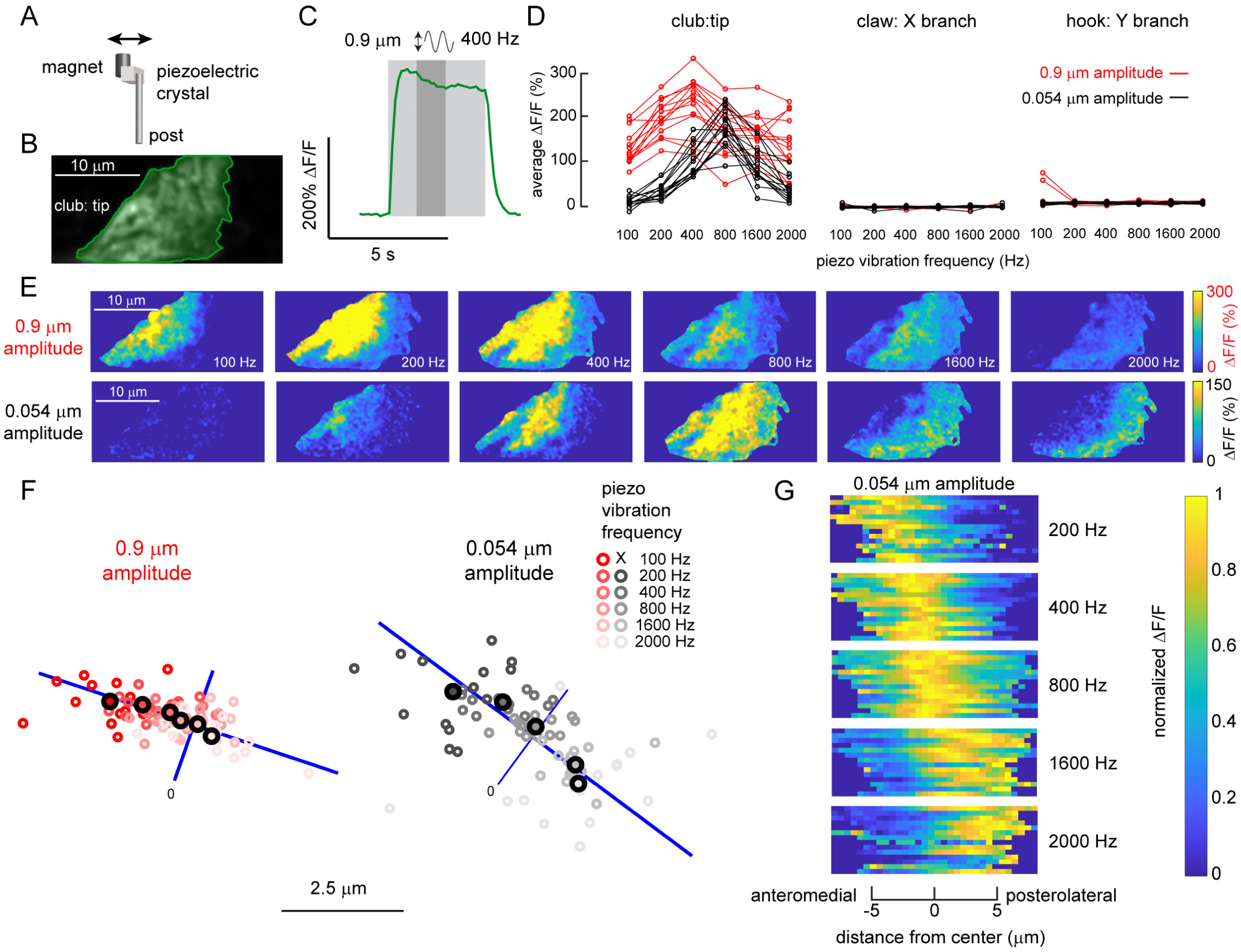
A map of vibration frequency in the axons of club neurons. **A.** We attached one side of a piezoelectric crystal to a magnet and the other to a post fixed to a servo motor. The magnet was placed directly onto a pin glued to the tibia (see Supplemental Figure 4 for details). **B.** Average GCaMP6f fluorescence from an example recording location, at the tip of the club axons (*R64C04-Gal4*). **C.** An example time course of the club’s response (ΔF/F) to a 400 Hz, 0.9 μm vibration of the magnet. We averaged the activity level in a 1.25 second window (indicated by a darker grey shading) starting from 1.25 seconds after the stimulation onset, and used it as a measure of response amplitude in D and as activity maps in E. **D.** Only club neurons respond reliably to the vibration stimulus. Plots show the activity of different subsets of FeCO axons in response to tibia vibrations at different frequencies and amplitudes. Each line represents an average response from one fly. **E.** Example ΔF/F maps of GCaMP6f fluorescence at the tip of the club in response to the vibration of the tibia at different frequencies and amplitudes. For both amplitudes, the responding regions shifted from the anterolateral side to the posteromedial side of the axon projection as stimulation frequency increased. **F.** A map of vibration frequency in club axon terminals. Smaller empty circles with different shades of red (0.9 μm amplitude) or grey (0.054 μm amplitude) represent the location of the weighted center of the responding region in different flies (n = 14 flies for both amplitudes). Larger, outlined circles represent the average location. The blue line represents the best fit line to the average locations across frequencies. We rotated the images from each fly to match the orientation of the example images shown in E. **G.** Distribution of activity (ΔF/F) along the anterolateral to posteromedial axis (blue lines in F) of the club axons during tibia vibration. Signals were normalized by the maximum average activity during each stimulus in that fly. Responses shifted from the anterolateral to posteromedial side as vibration frequency increased.

Hook neurons phasically increased their activity during tibia flexion but not extension (**Figure 5C**). Similar to the club neurons, hook responses were slightly weaker at fully extended positions. Phasic responses to tibia movement decayed rapidly, and we never observed tonic activity during the hold period. Overall, we found that hook neurons are directionally-selective, and encode flexion movements of the tibia.

To analyze the position-dependence of claw neurons in more detail, we measured steady state calcium activity at the end of each hold period (**Figure 5D, inset**) and plotted it against the femur-tibia angle (**Figure 5D**). To compare activity across flies, we normalized the response amplitudes from each fly with the largest steady state response recorded in that fly. The position tuning of claw neurons was relatively consistent across flies (**Figure 5D**) and branches of the claw. However, we did observe differences in claw position tuning across stimuli. Specifically, steady state activity was larger when the tibia was moved in the direction that increased claw activity (**Figure 5D-E**). This phenomenon, commonly referred to as hysteresis, would introduce ambiguity for downstream neurons trying to decode absolute leg angle. We did not observe directional hysteresis in the club or hook neurons.

For movement-sensitive club and hook neurons, we also examined velocity tuning using swing stimuli across a range of speeds (100-800 °/s). Because the amplitude of calcium signals reflects the integration of voltage changes over time, we compared the maximum slope of the calcium signals at different tibia movement speeds (**Figure S2D, E**). For club neurons, the slope of the calcium signal peaked around 400 °/s and decayed slightly at higher speeds (**Figure S2D**). In contrast, the slope of the calcium signal in hook neurons was similar across the entire speed range we tested (**Figure S2E**). These results suggest that in addition to their directional selectivity, club and hook neurons may also differ slightly in their velocity sensitivity.

### Club axons respond to low amplitude vibrations of the tibia and contain a spatial map of frequency

While imaging from club neurons, we occasionally observed sustained bursts of axonal calcium following a ramp movement of the tibia. We hypothesized that these variable signals were caused by spontaneous leg movements below the spatial resolution of our imaging system (3.85 μm/ pixel). Indeed, previous recordings in other insects have shown that FeCO neurons can be sensitive to low amplitude vibrations (Field and Pflüger, 1989; Stein and Sauer, 1999). To test this hypothesis, we used a piezoelectric chip to vibrate the magnet in a sinusoidal pattern (peak-to-peak amplitude 0.9 μm or 0.054 μm) at different frequencies (100 to 2000 Hz) (**Figure 6A**) and recorded calcium signals from club, claw, and hook neurons (see **Figure S3** for calibration details).

Club neurons exhibited large, sustained calcium signals to tibia vibration (**Figure 6**). An example trace in **Figure 6C** shows the response of club axons to 4 seconds of a 400 Hz vibration stimulus. Claw and club neurons did not respond to these stimuli (**Figure 6D**). To examine frequency tuning of club neurons, we averaged the calcium signal across the stimulus period (darkly shaded region in **Figure 6C**) and compared responses across different frequencies and amplitudes for multiple flies (**Figure 6D**). For the larger amplitude stimulus (0.9 μm), the club neurons showed significant responses at all frequencies, peaking at 400 Hz (**Figure 6D, red**). The responses to the smaller amplitude vibration stimulus (0.054 μm) were slightly weaker and peaked at 800 Hz (**Figure 6D, black**).

When we examined the distribution of calcium signals during tibia vibration, we noticed a consistent spatial shift in activity as a function of vibration frequency (**Figure 5E**). Specifically, the center of the calcium response moved from anterolateral to the posteromedial side of the club axon projection as frequency increased (**Figure 6F**). This shift was clearly visible in ΔF/F maps of the club axons for both large (0.9 μm) and small (0.054 μm) amplitude vibrations (**Figure 5E**). The frequency map was also consistent across flies (**Figure 6F-G**). Although the magnitude of the shift was larger for smaller vibrations, the shift was statistically significant for both vibration amplitudes.

The spatial maps in **Figure 6F** are not merely due to additional axons being recruited at higher frequencies. When we plotted the average activity along the anterolateral to posteromedial axis of the club axon projection for each fly, we found a significant shift in the entire response distribution as vibration frequency increased (**Figure 6G**). This effect was larger for the smaller vibration stimulus (0.054 μm), perhaps due to the saturation of calcium signals during larger amplitude vibration (0.9 μm). Overall, our imaging data reveal the existence of a frequency map within the axon terminals of club proprioceptors.

## Discussion

In this study, we used *in vivo* calcium imaging to investigate the population coding of leg proprioception in the femoral chordotonal organ (FeCO) of *Drosophila*. Our results reveal a basic logic for proprioceptive sensory coding: genetically distinct proprioceptor subtypes detect and encode distinct kinematic features, including tibia position, directional movement, and vibration. The cell bodies of each proprioceptor subtype reside in separate parts of the FeCO in the leg, and their axons project to distinct regions of the fly ventral nerve cord (VNC). This organization suggests that different kinematic features may be processed by separate downstream circuits, and function as parallel feedback channels for the neural control of leg movement and behavior.

### Neural representation of tibia position

Claw neurons encode the position of the tibia relative to the femur. Specifically, each branch consists of two sub-branches, whose calcium signals increase when the tibia is flexed or extended (**Figure 5**). When the tibia is close to 90°, both sub-branches are minimally active. This response distribution is similar to that observed in multiunit recordings from the FeCO of locusts and stick insects (Burns, 1974; Usherwood et al., 1968). However, single unit recordings from these species revealed the existence of a small number of position-tuned cells with peak activity in this middle range (Hofmann et al., 1985; Matheson, 1990; Matheson, 1992). We did not find such responses in our calcium imaging experiments. Because we imaged from multiple neurons simultaneously, we could have missed weak signals from a small number of axons. It is also possible that the driver lines we used did not label the FeCO neurons tuned to this range (**Figure 3**). Finally, it is possible that this represents a real difference between *Drosophila* and other insects. The fly FeCO has about half as many neurons as that of the stick insect and locust, and the biomechanics of the organ may also differ between species (Field, 1991; Shanbhag et al., 1992). Future experiments that characterize the position-tuning of single neurons in the fly may help to distinguish between these possibilities.

Position-encoding claw neurons exhibit response hysteresis: both flexion and extension-tuned sub-branches of the claw showed larger steady state activity when the tibia is moved in a direction that increases their activity (**Figure 5**). This response asymmetry is notable because it presents a problem for downstream circuits and computations that rely on a stable readout of tibia angle. Proprioceptive hysteresis has also been described in vertebrate muscle spindles (Yu Wei et al., 1986) and FeCO neurons of other insects (Matheson, 1992). One possible solution for solving the ambiguities created by hysteresis would be to combine the tonic activity of claw neurons with signals from directionally selective hook neurons (**Figures 4, 5**). This could allow a neuron to decode tibia position based on past history of tibia movement. However, it is also possible that tibia angle hysteresis is a useful feature of the proprioceptive system, rather than a bug. For example, it has been proposed that hysteresis could compensate for the nonlinear properties of muscle activation in short sensorimotor loops (Zill and Jepson-Innes, 1988).

### Neural representation of tibia movement/vibration

We identified two functional subtypes of FeCO neurons that respond phasically to tibia movement. Club neurons respond to both flexion and the extension of the tibia, while hook neurons respond only to flexion (**Figures 4, 5**). In population-imaging experiments, we also observed directionally-tuned responses selective for tibia extension (**Figure 2**). Overall, the movement sensitivity of the club and hook neurons resembles that of other phasic proprioceptors, including primary muscle spindle afferents (Jones et al., 2001), and movement-tuned FeCO neurons recorded in the locust and stick insect (Hofmann et al., 1985; Matheson, 1990; Matheson, 1992). Although the slow temporal dynamics of GCaMP did not permit a detailed analysis of velocity tuning, our results indicate that FeCO neurons respond to the natural range of leg speeds encountered during walking (Mendes et al., 2013; Wosnitza et al., 2013).

In addition to their directional tuning, we found that club and hook neurons differ in their sensitivity to fast (100 Hz – 2000 Hz), low amplitude (0.9 μm – 0.054 μm) tibia vibration. Club neurons are strongly activated by vibrational stimuli, but hook neurons are not (**Figure 6**). This difference in vibrational sensitivity is not likely to be caused by a difference in velocity tuning, because the subtypes respond similarly to the range of the speeds experienced during tibia vibration (**Figure S2**). Rather, it seems that the club neurons have a lower mechanical threshold, and/or may be more sensitive to the constant acceleration produced by vibration.

Although club neurons account for ∼30% of the FeCO neurons in *Drosophila*, and other insects also possess large numbers of vibration-sensitive FeCO neurons (Field and Pflüger, 1989; Stein and Sauer, 1999), their functional role is still not clear. Previous studies in stick insects and locusts have found that vibration-tuned FeCO neurons do not contribute to postural reflexes like other FeCO neurons (Field and Pflüger, 1989; Kittmann and Schmitz, 1992; Stein and Sauer, 1999). One possibility is that club neurons monitor substrate vibrations, which may be important for fly courtship (Fabre et al., 2012) and other forms of intraspecific communication (Mazzoni et al., 2013). However, another possibility is that vibration sensitivity is an artifact of mechanical tuning for higher-order kinematic parameters, such as joint acceleration (Hofmann and Koch, 1985). An analogous example is that of muscle spindles, in which tendon vibration drives 1a afferent activity and produces illusions of limb movement (Roll et al., 1989). Such vibration stimuli may rarely occur in a natural setting, and so may not be ethologically relevant. Having genetic access to proprioceptor subtypes in the fly should facilitate a functional dissection of how vibration sensors contribute to motor control and behavior.

### Stereotypic spatial organization of leg proprioceptors

Using genetic driver lines for specific FeCO neuron subtypes, we provide the first detailed anatomical characterization of *Drosophila* leg proprioceptors. Our anatomy and imaging experiments revealed a systematic relationship between the functional tuning of proprioceptor subtypes and their anatomical structure. The cell bodies of the three proprioceptor subtypes are clustered in different regions of the femur, an organization that may reflect biomechanical specialization for detecting position, movement, and vibration. Proprioceptor axons then converge within the leg nerve, before branching within the VNC to form subtype-specific projections that we call the club, claw, and hook (**Figures 2, 3, and 4**). We found that this organization is highly stereotyped across flies.

The axons of claw neurons split into three symmetric branches, resembling a claw. This unique arborization pattern is suggestive of a Cartesian coordinate system; for example, each branch could represent a different spatial axis. However, we found that each claw neuron innervates all three branches, and that the X, Y, and Z branches all encode the same stimuli. Specifically, our calcium imaging experiments revealed that each claw branch is divided into two sub-branches that are specialized for encoding flexion or extension of the tibia (**Figures 2, 4**). If each claw branch is functionally similar, what is the purpose of this tri-partite structure? Each branch may target different downstream neurons, or could be independently modulated by presynaptic inhibition. Interestingly, the axons of directionally-tuned hook neurons arborized alongside the claw, but did not innervate all three of the claw branches. Thus, the X, Y, and Z branches may facilitate integration of positional information with directionally-tuned movement signals.

We were surprised to discover a topographic map of leg vibration frequency within the axon terminals of club neurons. This structure has not previously been described in flies, but resembles the tonotopic map of sensory afferents in the cricket auditory system (Oldfield, 1983; Romer, 1983) or the cochlear nucleus in vertebrates (Cohen and Knudsen, 1999). Interestingly, the spatial layout of the frequency map in club axons was consistent across different vibration amplitudes, despite a shift in the peak frequency tuning curve (**Figure 6**). An orderly map of vibration frequency could facilitate feature identification in downstream circuits, for example through lateral inhibition between neighboring axons with shared tuning (Suga, 1989).

### Comparison with other sensory systems

Neurons in the FeCO population can be generally classified as either tonic (position-encoding) or phasic (movement-encoding). This division has been observed in the proprioceptive systems of many animals, including other insects (Hofmann et al., 1985; Matheson, 1990; Zill, 1985), crustaceans (Burke, 1954), and mammals (Proske and Gandevia,2012). For example, mammalian muscle spindles are innervated by both phasic (Group 1a) and tonic (Group II) afferents (Boyd, 1980). The same is true of other mechanosensory modalities, including touch (Abraira and Ginty, 2013; Burrows and Newland, 1997; Juusola and French, 1998) and hearing (Kamikouchi et al., 2009; Yorozu et al., 2009). The ubiquity of tonic and phasic sensory neurons suggests that these two parallel information channels are essential building blocks of sensory circuits. Now that we have identified genetic tools that delineate tonic and phasic neurons in the proprioceptive system of *Drosophila*, this system has the potential to provide general insights into the utility of this sensory coding strategy.

## Summary

With the advent of new methods for simultaneously monitoring the activity of hundreds or thousands of neurons (Ahrens et al., 2013; Mann et al. 2017; Sofroniew et al., 2016), a critical challenge has been to link the activity of large neuronal populations to the underlying diversity of specific cell-types (Alivisatos et al., 2013). Previous efforts have used statistical methods to compare the responses of single neurons to simultaneous optical (Tsodyks et al., 1999) or electrophysiological (Okun et al., 2015) population recordings. Here, we took a different approach, which took advantage of the fact that neurons in the fly can be reliably identified across individuals. We first used 2-photon imaging to monitor activity across a population of proprioceptive sensory neurons during controlled leg movements. From this population data, we identified spatially distinct axon branches that encode specific proprioceptive stimulus features. We then searched for genetic driver lines that specifically labeled each axon branch, and further characterized their functional tuning with targeted calcium imaging. With this approach, we were able to identify and characterize the major neuronal subtypes in a key proprioceptive organ.

With a genetic handle on position, movement, and direction pathways, it should now be possible to trace the flow of proprioceptive signals into downstream circuits, and to identify the functional role of specific proprioceptor subtypes within the broader context of motor control and behavior. We anticipate that *Drosophila* will provide a useful complement to other model organisms in dissecting fundamental mechanisms of proprioception, and deepening our understanding of this mysterious “sixth sense”.

## Acknowledgements

We thank Eiman Azim, Richard Mann, Jim Truman, Julijana Gjorgjieva, and members of the Tuthill laboratory for helpful comments on the manuscript. This work was funded by a Searle Scholar Award, a Klingenstein-Simons Fellowship, and NIH grant R01NS102333 to J.C.T.

## Author Contributions

A.M. and P.G. performed the experiments, A.M., P.G., and J.C.T. analyzed the data, and A.M. and J.C.T. designed the experiments and wrote the manuscript.

## Methods

### Fly preparation for in vivo two-photon calcium imaging of FeCO axons

*Drosophila melanogaster* were raised on a standard cornmeal medium and kept at 25°C in a 12:12 h light dark cycle. We used female flies 1 to 4 days post-eclosion for all experiments. To gain optical access to the VNC while moving the tibia, we used a previously described fly holder (Tuthill and Wilson, 2016) (Figure 1B), but replaced the metal sheet that holds the fly’s thorax with a thin, translucent plastic sheet. The plastic sheet served as a light diffuser and provided a bright background during the automated tracking of the tibia. To place the fly into the holder, we first anesthetized the fly by cooling in a plastic tube on ice, then put the fly’s head through the hole in the holder and glued the ventral side of the thorax onto the hole using UV-cured glue (Bondic, Aurora, ON, Canada). We glued the head to the upper side of the fly holder. On the bottom side of the holder, we glued down the femur section of the right prothoracic leg so that we could control the femur-tibia joint angle by moving the tibia. When gluing the femur, we held it at a position where the movement of the tibia during the rotation of the femur-tibia joint was parallel to the plane of the fly holder. To eliminate mechanical interference, we glued down all other legs. We also pushed the abdomen to the left side and glued it at that position, so that the abdomen did not block tibia movement during flexion. To position the tibia with the magnetic control system described below, we cut a small piece of insect pin (length ∼1.0 mm, 0.1 mm diameter; Living Systems Instrumentation, St Albans, VT) and glued it onto the tibia and the tarsus of the right prothoracic leg. We painted the pin with black India ink (Super Black, Speedball Art Products, Statesville, NC) to enhance contrast and improve tracking of the tibia/pin position. After immersing the preparation in *Drosophila* saline, we removed the cuticle above the prothoracic segment of the VNC with fine forceps and took out the digestive tract to reduce the movements of the VNC. We also removed fat bodies and larger trachea to improve the optical access to the leg neuropil. Fly saline contained: 103 mM NaCl, 3 mM KCl, 5 mM TES, 8 mM trehalose, 10 mM glucose, 26 mM NaHCO3, 1 mM NaH2PO4, 1.5 mM CaCl2, and 4 mM MgCl2 (pH 7.1, osmolality adjusted to 270-275 mOsm).

Recordings were performed at room temperature.

### Image acquisition using a two-photon excitation microscope

We used a modified version of a custom two-photon microscope previously described in detail (Euler et al., 2009). For the excitation source, we used a mode-locked Ti/sapphire laser (Mira 900-F, Coherent Inc, Santa Clara, CA) set at 930 nm and adjusted the laser power using a neutral density filter to keep the power at the back focal plane of the objective (40x, 0.8 NA, 2.0 mm wd; Nikon Instruments Inc, Melville, NY) below ∼25 mW during the experiment. We controlled the galvo laser scanning mirrors and the image acquisition using ScanImage software (version 5.2) running on Matlab (MathWorks, Natick, MA). To detect GCaMP6f and Td-Tomato fluorescence, we used a ET510/80M (Chroma Technology Corporation, Bellows Falls, VT) emission filter (GCaMP6f) or a 630 AF50/25R (Omega optical Inc, Brattleboro, VT) emission filter (Td-Tomato) and GaAsP photomultiplier tubes (H7422P-40 modified version without cooling; Hamamatsu Photonics, Japan). During the trials, we acquired images at 8.01 Hz. At the end of the experiment, we acquired a z-stack of the FeCO axons in the right hemisphere of the prothoracic leg neuropil to confirm the recording location.

### Moving the tibia/pin using a magnetic control system

To move the tibia/pin to different positions, we attached a rare earth magnet (1 cm height x 5 mm diameter column) to a steel post (M3×20mm flat head machine screw) and controlled its position using a computer programmable servo motor (SilverMax QCI-X23C-1; Max speed 24,000 deg/s, Position resolution 0.045 deg; QuickSilver Controls, Inc, San Dimas, CA). We placed a piezoelectric crystal (PA3JEW; ThorLabs, Newton, NJ) between the post and the magnet in order to vibrate the tibia as described below. To move the magnet in a circular trajectory centered at the femur-tibia joint, we placed the motor on a micromanipulator (MP-285, Sutter Instruments, Novato, CA) and adjusted its position while visually inspecting the movement of the magnet and the tibia using the tibia tracking camera described below. This brought the top edge of the magnet to the same height as the tibia/pin, and the inner edge of the magnet to be 1.5 mm from the center of the femur-tibia joint (Figure 1C). Because the pin was glued slightly off the center of the joint, the distance between the pin and the magnet was approximately 300 μm. For each trial, we controlled the speed and the position of the servo motor using QuickControl software (QuickSilver Controls, Inc).During all trials, we tracked the tibia position (as described below) to confirm the tibia movement during each trial.

Because it was difficult to fully flex the femur-tibia joint without the tibia/pin and the magnet colliding with the abdomen, we only flexed the joint up to ∼18°. During the swing motion trials (Figures 2 and 4), we commanded the motor to move from a fully extended position (180°) to a fully flexed position (18°) at 360 °/s and move back to the original position with the same speed. (For velocity sensitivity experiments in Figure S2D-E, we also used 180, 720, and 1440 °/s movements.) There was a 5-second interval between the flexion and the extension movement. We repeated this swing motion 3 times with a 5-second inter-trial interval and averaged the responses within each fly prior to averaging across flies.

For ramp-and-hold motion trials (Figure 5), we programmed the motor to move in 18° steps between full extension and flexion at 240 °/s. There was a 3-second hold step between each ramp movement. We used two types of movements for a ramp-and-hold motion: one that started with a flexion of the joint and extended back to the original position and another that started with an extension of the joint and flexed back to the original position. We repeated each type of trial twice and averaged the responses within the preparation before averaging them across preparations. We set the acceleration of the motor to 72000 °/s^2^ for all movements. Movements of the tibia during each trial varied slightly due to several factors, including a small offset between the center of the motor rotation and the femur-tibia joint, and the acceleration and deceleration of the tibia movement in response to the magnet motion. Because these variations were relatively small (judging from how the responses changed with speed in Figure S2D-E), we did not consider these differences in the initial summary of the responses to swing motion and ramp-and-hold motion (Figure 2, 4, and 5A-C). However, for quantifying the positional dependence of the responses, we plotted the response against the actual position of the tibia during each trial (Figure 5D). Because the actual position of the tibia differed slightly between preparations, we interpolated the data at 5° intervals when averaging position dependent responses across preparations.

### Tracking the femur-tibia joint angle

To track the position of the tibia, we backlit the tibia/pin with an 850 nm IR LED (M850F2, ThorLabs) and recorded video using an IR sensitive high speed video camera (Basler Ace A800-510um, Basler AG, Ahrensburg, Germany) with a 1.0x InfiniStix lens (94 mm wd, Infinity, Boulder, CO) equipped with 900 nm short pass filter (Edmund optics, Barrington, NJ) to filter out the two-photon laser light (Figure 1B). Because the servo motor was directly underneath the fly, we placed the camera on the side and used a prism to capture the view from below. We recorded the images at 180 Hz for the ramp-and-hold motion, and at 200 Hz for the swing motion (exposure time 2.5 ms for both types of motion). To synchronize the images taken by the camera with those taken by the two-photon microscope, we acquired both the camera exposure signal and the position of the galvo scanning mirrors at 20 kHz during the trials. After acquiring the images (Figure 1C), we identified the position of the dark tibia/pin against the bright background by thresholding the image. We then approximated the orientation of the leg as the long axis of an ellipse with the same normalized second central moments as the thresholded image (Haralick and Shapiro, 1992). The spatial resolution of the image was 3.85 μm per pixel and (assuming circular movement of the tibia/pin), 1-pixel movement at the edge of the tibia/pin (∼1.2 mm from the center of the rotation) corresponded to 0.18°.

### Image processing, K-means clustering of the responses, and analyses of clustered responses

We performed all image processing and analyses using scripts written in Matlab (MathWorks). After acquiring the images for a trial, we first applied a Gaussian filter (size 5×5 pixel, σ = 3) and aligned each frame to a mean image of the trial using a sub-pixel registration algorithm (registered to 1/4 pixel) (Guizar-Sicairos et al., 2008). For alignment of images, we used Td-tomato fluorescence. For detecting calcium signals, we chose pixels whose mean GCaMP6f fluorescence was above a set threshold. We used correlation based K-means clustering to group pixels based on the similarity of their intensity change during the trial (Figure 1F). We initially chose the number of clusters based on visual inspection of the response patterns during individual trials. After this initial clustering, we manually adjusted the number of clusters to verify response separability.

For calculating the GCaMP6f fluorescence change relative to the baseline (ΔF/F), we used the lowest average fluorescence level in a 10-frame window as the baseline fluorescence during that trial. To group similarly-responding clusters across flies in an unbiased fashion, we collected the ΔF/F traces recorded from each region in all flies and performed another correlation based K-means clustering. As in the clustering for each trial, we initially chose the number of groups based on the visual inspection of the responses recorded from that region. We varied the group numbers for each region and selected the number of groups that best categorized the response patterns we observed. The results of this second clustering exercise are indicated by the red, blue, green, purple, and orange response types in Figure 2.

When comparing the time course of the same type of responses recorded from different axonal regions during swing motion, we first normalized each response time course (74 frames starting from 10 frames before the onset of the flexion) with the maximum amplitude for that response. After the normalization, we calculated the root mean square error for each response time course against the average response time course for that region, or for all the recording regions (Figure S1B-C, S2A-C). If the root mean square errors in both cases were not statistically different (t-test), we considered that the time course of the same response type recorded from different regions were not statistically significantly different.

To investigate the velocity sensitivity of the club and hook neurons, we first calculated the maximum slope of the ΔF/F curves for these neurons during swing motion (both flexion and extension for club neurons, flexion only for hook neurons). We reasoned that the maximum slope of the ΔF/F curves more accurately represents the maximum activity level of these neurons than the maximum amplitude of the ΔF/F curves, because the calcium signal integrates the activity of the neuron over time. For each response, we calculated the average tibia speed during the inter-frame interval that showed the maximum slope, and plotted the maximum slope against this speed (Figure S2D, E).

### Analysis of the spatial organization of each response type

After identifying different response types for each trial, we inspected the spatial organization of pixels that belonged to each response type. Because these pixels were grouped together in space, we used the center of mass of these pixels as a measure of their location and calculated their position relative to the center of mass of all the GCaMP6f positive pixels in the image. We used the characteristic shapes of the axon projections as landmarks and recorded from similar locations in different flies, but the anterior-posterior axis of the VNC in the image slightly differed among the preparations (Figure 2A-D, left columns). Before analyzing the spatial organization of each response types, we compensated for these differences by rotating the images. For each recording region, we rotated the images from different flies to align the orientation of the claw axon projection (for recordings near the X and Y branches), the anterior edge of the club axon projection (for recordings near the tip of the club), or the axis connecting the Z branch and the directionally selective branch (for recordings near the Z branch) in the images (Figure 2A-D, right columns). Figure S1A shows outlines of the GCaMP6f positive pixels from different *iav-Gal4 x UAS-GCaMP6f* flies for each recording location after these compensations. As can be seen from the Figure, these rotations were able to account for most of the differences in the recording region orientations. For a statistical comparison of the locations of the different response types in *iav-Gal4 x UAS-GCamp6f* flies, we calculated the relative orientation and distance between each cluster’s center of mass and constructed a vector plot (Figure S1D-G). For each cluster pair, we first calculated a mean vector and took each vector’s component along the mean vector orientation. Then, we ran one-tailed Wilcoxon signed rank test on those components to test if they were statistically significantly positive. If they were significantly positive, we considered the centers of mass for these two clusters to be statistically different.

### Vibrating the tibia using a piezoelectric crystal

To vibrate the tibia at high frequencies, we moved the magnet using a piezoelectric crystal (PA3JEW, Max displacement 1.8 μm; ThorLabs) (Figure 6A). For controlling the movement of the piezo, we generated sine waves of different frequencies in Matlab (sampling frequency 10 kHz) and sent them to the piezo through a single channel open-loop piezo controller (Thorlabs). Because tibia movements induced by the piezo electric crystal were below the resolution of our tibia tracking system (3.85 μm/pixel), we first calibrated the piezo induced tibia movements using a separate tracking system equipped with a long working distance high magnification objective (50x, 0.45 NA, Nikon) connected to a high speed video camera (A800-510um, Basler) via InfiniTube FM-100 (Infinity) (Figure S3). In this setup, we were able to record images at 4000 Hz (exposure 190 μs) with a spatial resolution of 0.106 μm/pixel. For better control of the tibia position during high-frequency vibration, we attached the magnet to the pin for all vibration experiments (magnet overlapped with the pin for ∼200 μm). For each vibration frequency, we measured the amplitude of the tibia oscillation envelope (Figure S3B, D) and power spectrum of the tibia movement (Figure S3C, E; Thomson’s multitaper power spectral density estimate with time-half bandwidth product = 4). The power spectrum of tibia movement showed a large peak at the command frequency of the piezo electric crystal, suggesting that most of the vibrations were indeed at the target frequency (Figure S3C). Both the amplitude of the oscillation envelope and the power of the oscillation at the target frequency decreased greatly at around 2000 Hz (Figure S3D, E). Thus, we decided to use vibration stimuli up to 2000 Hz.

For each stimulus, we presented 4 seconds of vibration twice with an inter-stimulus interval of 8 seconds. We averaged the responses within each fly before averaging across flies. For calculating ΔF/F maps and average response amplitudes, we used the activity level in a 1.25-second window starting 1.25 seconds after the vibration onset.

### Analyzing frequency tuning map within the club axon projection

Because the orientation of the club axon projection in the recorded image differed slightly between preparations, we first rotated the images so that the orientation of the anterior edge of the club axon projection matched the example images in Figure 6E. In each fly, we defined the center of the response for each vibration frequency as the weighted center of mass of the average ΔF/F map for each stimulus. To see how the location of the response centers changed with the vibration frequency, we plotted the response centers relative to the center of mass for all the GCaMP6f positive pixels in the image (Figure 6F). After calculating the location of the average response centers for each vibration frequency, we fit a line through them by taking the first principal component of the x, y coordinates of the average response centers as the slope of the line. This line minimizes the total distance between the line and average response centers. To calculate the response distribution along this line (Figure 6G), we binned (5 pixels/bin) the pixels based on the distance along this line and averaged their ΔF/F values. For each vibration frequency in each fly, we normalized the response distribution by the maximum response.

### Immunohistochemistry and anatomy

For confocal imaging of the FeCO neuron axons driven by each Gal4 line in the VNC (Figure 3A), we crossed flies carrying the Gal4 driver to flies carrying *pJFRC7-20XUAS-IVS-mCD8::GFP* and dissected the progeny’s VNC out of the thorax in a *Drosophila* saline. We first fixed the VNC in a 4% formaldehyde PBS solution for 15 minutes. After rinsing the VNC in PBS three times, we put it in the blocking solution (5% normal goat serum in PBS with 0.2% Trition-X) for 20 minutes, then incubated it with a solution of primary antibody (anti-GFP chicken antibody 1:50 concentration; anti-brp mouse for neuropil staining; 1:30 concentration) in blocking solution for 24 hours at room temperature. At the end of the first incubation, we washed the VNC in PBS with 0.2% Triton-X (PBST) three times, then incubated the VNC in a solution of secondary antibody (anti-chicken-Alexa568 1:250 concentration; anti-mouse-Alexa488 1:250 concentration) dissolved in blocking solution for 24 hours at room temperature. Finally, we washed the VNC in PBST for three times and then mounted it on a coverslip with Vectashield (Vector Laboratories, Burlingame, CA). We acquired a z-stack image of the slides on a confocal microscope (FluoView 1000; Olympus, Center Valley, PA) to capture the axonal projection pattern relative to the neuropil.

For multicolor flip-out experiments (Nern et al., 2015; Figure 3B), we crossed flies carrying the multicolor flip out cassettes and Flp recombinanse drivers to flies carrying different Gal4 drivers, and dissected the progeny. For temperature induced expression of Flp, we placed adult flies in a plastic tube and incubated them in a 37°C water bath for 13 to 15 minutes (up to 1 hour for the *R21D12-Gal4* flies). We dissected the VNC 3 days after the Flp induction and followed the procedure described above to detect HA (using anti-HA-rabbit antibody), V5 (using DyLight 549-conjugated anti-V5), and FLAG (using anti-FLAG-rat antibody) labels expressed due to the Flp induction in individual neurons.

For co-labeling of the cell bodies of FeCO neurons driven by each Gal4 line with those of all FeCO neurons (driven by ChaT-LexA) (Figure 3C-F), we crossed flies carrying the UAS-Red-Stinger and LexAop-GFP with flies carrying one of the Gal4 driver and Chat-LexA. We used the progenies of this cross that express red-stinger in the nucleus of FeCO neurons driven by the Gal4 line and GFP in the nucleus of all FeCO neurons. For immunostaining of the leg, we followed the same procedure as those described above for the VNC except that we incubated the leg longer in the antibodies to allow the antibodies to penetrate in to the leg.

For *in silico* overlay of the expression patterns of specific Gal4 lines (Figure 3G), we used confocal stacks of each Gal4 line with neuropil counterstaining (from the Janelia FlyLight database (Jenett et al., 2012)) and used the neuropil staining to align the expression pattern in the VNC using the Computational Morphometry Toolkit (CMTK; Jefferis et al., 2007).

---

**Figure S1:**
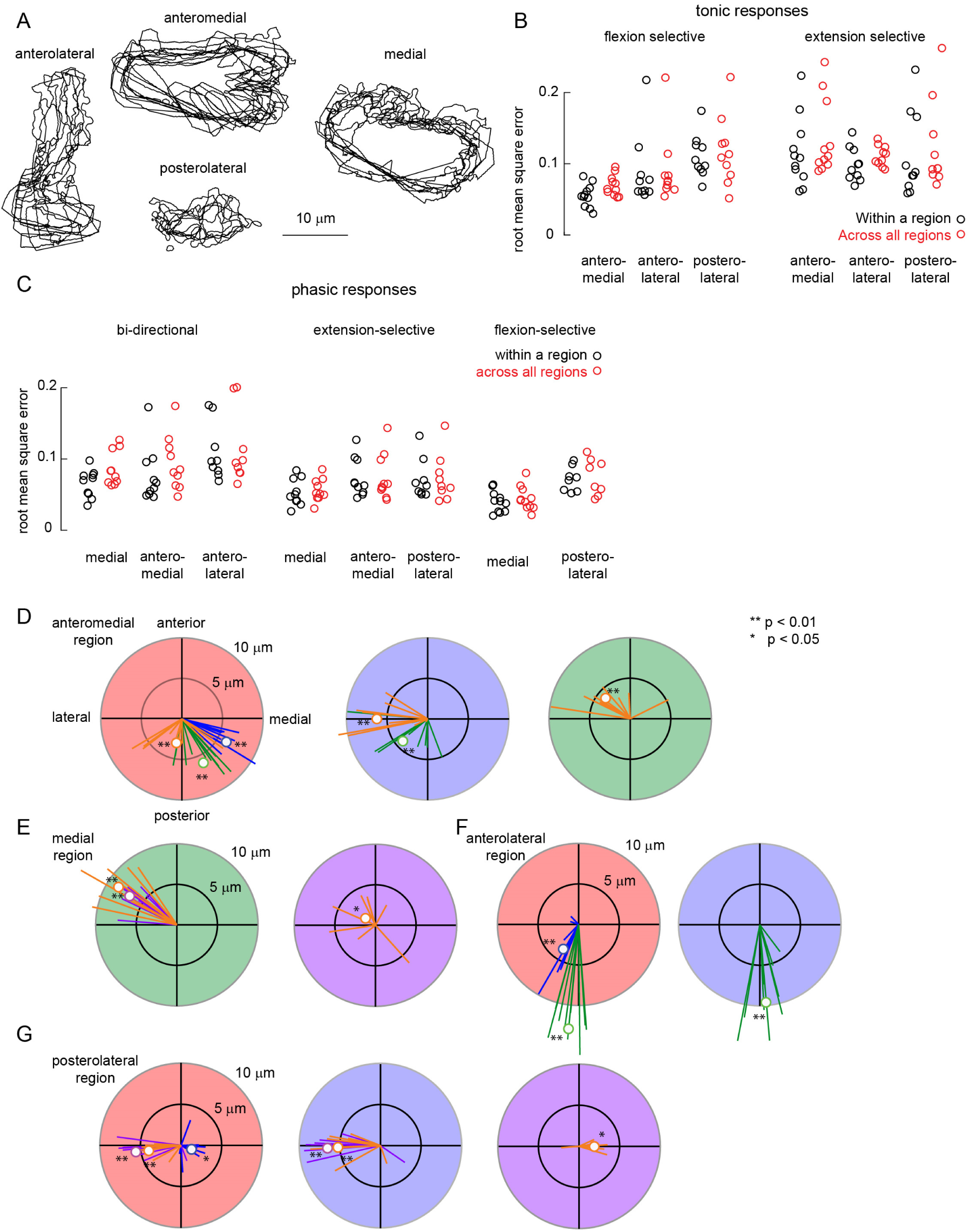
related to Figure 2. Spatial and temporal properties of population responses from *FeCO neurons.* **A.** Outlines of pixels containing GCaMP6f fluorescence in recordings at each location from different flies (summary of responses shown in **Figure** 2A-D). In order to align recordings from different flies, we rotated the images in each fly as described in the **Figure** 2A-D legends. Anterolateral, n = 10; anteromedial, n= 11; postero-lateral, n = 10, medial, n = 11. **B.** To compare response tuning across different axon projections, we computed the root mean square error (RMSE) of calcium signals from each fly to the average calcium signal across flies, both within individual regions (black) and across all regions of the same response type (red). The response time courses are summarized in **Figure** 2A-C. The RMSEs within and across regions were not statistically different. **C.** Same as B, but for phasic responses. Again, the RMSEs within and across regions were not statistically different. **D.** Polar plots showing the relative direction and distance between different response clusters recorded from the anteromedial region of iav-Gal4 (data correspond to the spatial layout plots in **Figure** 2B, and the color scheme is preserved). Polar plots shaded with light red, blue, and green, indicate the direction and distance from the pixels that showed flexion-selective tonic responses (n = 11), extension-selective tonic response (n= 11), and bi-directional phasic response (n= 10), respectively. Blue, green, and orange lines in each polar plot show the direction and distance to the pixels that showed extension-selective tonic responses, bidirectional phasic responses, and extension-selective phasic responses (n= 10), respectively. **E.** Same as D, but for responses recorded from the medial region (shown in **Figure** 2D). In addition to the response types described in D, we found flexion-selective phasic responses in this region, which are represented by purple lines and circles and a polar plot with light purple shading. Green: n = 10, orange: n =10, purple: n = 11. **F.** Same as D, but for responses recorded from the anterolateral region (shown in **Figure** 2A). Red: n = 10, blue: n = 10, green: n =9. **G.** Same as D, but for responses recorded from the posterolateral region (shown in **Figure** 2C). Red: n= 10, blue: n = 10, purple: n = 8, orange: n = 9.

**Figure S2:**
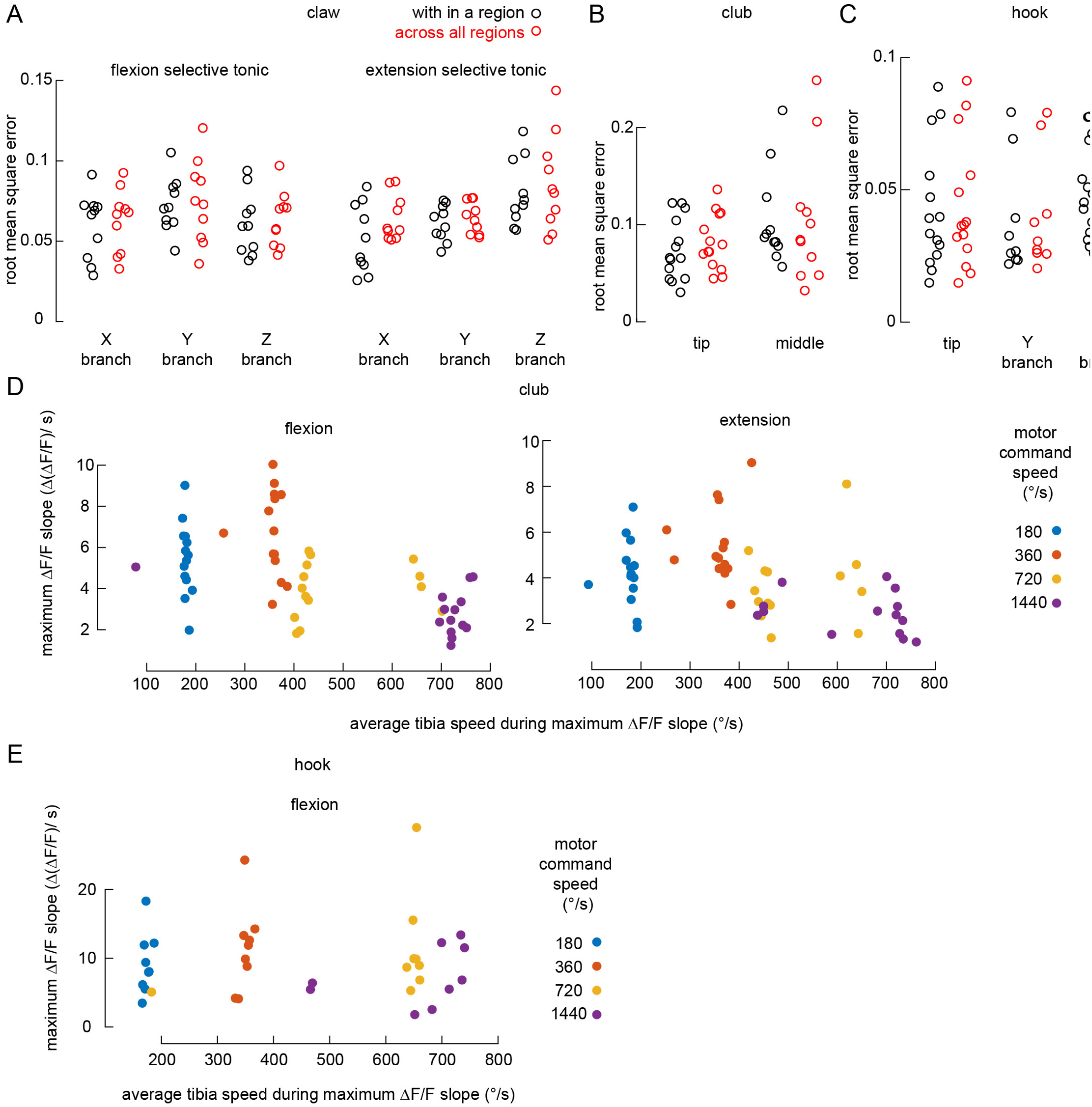
related to Figure 4. Spatial and temporal properties of subtype-specific responses. **A.** RMSEs for tonic responses recorded from each claw branch (*R73D10-Gal4*). For each fly, we compared the calcium signal of each region to the average response from the same region (black circles), or to the average of responses recorded from all regions (red circles). RMSEs within and across regions were not statistically different. Response time courses are shown in **Figure** 4A. **B.** Same as A, but for bidirectional phasic responses recorded from the tip (left) and the middle (right) of the club axon projection driven by *R64C04-Gal4*. RMSEs within and across regions were not statistically different. Response time courses are shown in **Figure** 4B. **C.** Same as A, but for flexion-selective phasic responses recorded from the tip, the Y branch, and the Z branch region of the hook axon projection driven by *R21D12-Gal4*. RMSEs within and across regions were not statistically different. Response time courses are shown in **Figure** 4C. **D.** Scatter plot showing the maximum slope of the ΔF/F curve for the club axons (recorded at the tip) as a function of average tibia swing speed. Each circle represents the average maximum slope of the ΔF/F curve in one fly. We calculated the average velocity of tibia movement from the frame interval in which we observed the maximum slope of the ΔF/F curve. For both flexion and the extension, n = 14 flies for all speeds. **E.** Same as D, but for hook axons recorded at the Y branch region. Because hook neurons respond only to the flexion, we only show the plot for flexion movements. n = 9 flies for all speeds.

**Figure S3:**
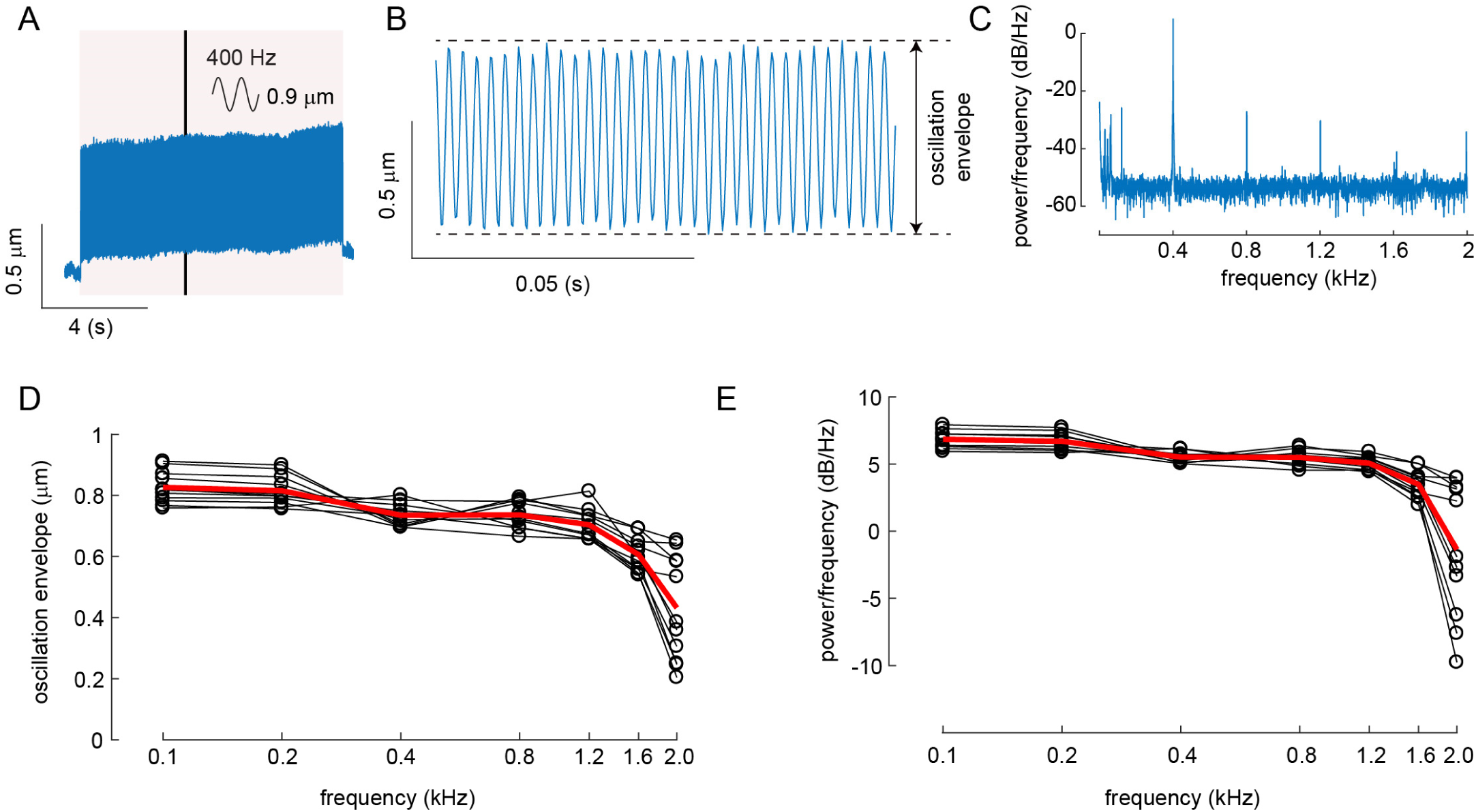
related to Figure 6. Calibration of vibration stimuli. **A.** An example tibia movement in response to a 400 Hz sine wave vibration of the piezoelectric crystal with 0.9 μm peak-to-peak amplitude. The grey shaded region indicates the duration of the piezo vibration. The black line represents the region expanded in B. **B.** An expanded view of the example tibia movement shown in A. **C.** The power spectrum of the example tibia movement shown in A. **D.** A plot showing how the oscillation envelope (as measured in B) of the tibia movement changed as we varied the command frequency of the piezoelectric crystal. Black lines indicate the oscillation envelope at different frequencies (n = 11 flies). The red line represents the mean oscillation envelope at different frequencies. The peak-to-peak amplitude of the piezo vibration was 0.9 μm for all frequencies. **E.** A plot showing how the power of the tibia vibration at the piezo command frequency changes as a function of command frequency. Black lines indicate the power of the tibia vibration at different frequencies for each fly (n = 11 flies). The red line represents the mean power of the tibia vibration at different frequencies.

## Table of genotypes

|  |  |
| --- | --- |
| **Figure 1A** | w[1118]; P{JFRC7-20XUAS-IVS-mCD8::GFP} attp40/+; Iav-Gal4/+ |
| **Figure 1D** | w[1118]; P{JFRC7-20XUAS-IVS-mCD8::GFP} attp40/+; Iav-Gal4/+ |
| **Figure 1E, G** | W[1118]/+; P{20xUAS-IVS-GCaMP6f} attP40 /+ ; Iav-Gal4/P{w[+mc]=UAS-tdTomato} |
| **Figure 2A-D** | W[1118]/+; P{20xUAS-IVS-GCaMP6f} attP40 /+ ; Iav-Gal4/P{w[+mc]=UAS-tdTomato} |
| **Figure 3 A** | w[1118]; P{JFRC7-20XUAS-IVS-mCD8::GFP} attp40/+; Iav-Gal4/+<br>w[1118]; P{JFRC7-20XUAS-IVS-mCD8::GFP} attp40/+; P{GMR73D10-GAL4}attP2/+<br>w[1118]; P{JFRC7-20XUAS-IVS-mCD8::GFP} attp40/+; P{GMR64C04-GAL4}attP2/+<br>w[1118]; P{JFRC7-20XUAS-IVS-mCD8::GFP} attp40/+; P{GMR21D12-GAL4}attP2/+ |
| **Figure 3B** | w[1118] P{y[+t7.7] w[+mC]=hs-FLPG5.PEST}attP3/ w[1118]; +/+; P{y[+t7.7]<br>w[+mC]=10xUAS(FRT.stop)myr::smGdP-OLLAS}attP2 PBac{y[+mDint2]<br>w[+mC]=10xUAS(FRT.stop)myr::smGdP-HA} VK00005 P{10xUAS(FRT.stop)myr::smGdP-V5-THS-10xUAS(FRT.stop)myr::smGdP-FLAG}su(Hw)attP1/ P{GMR73D10-GAL4}attP2<br>w[1118] P{y[+t7.7] w[+mC]=hs-FLPG5.PEST}attP3/ w[1118]; +/+; P{y[+t7.7]<br>w[+mC]=10xUAS(FRT.stop)myr::smGdP-OLLAS}attP2 PBac{y[+mDint2]<br>w[+mC]=10xUAS(FRT.stop)myr::smGdP-HA} VK00005 P{10xUAS(FRT.stop)myr::smGdP-V5-THS-10xUAS(FRT.stop)myr::smGdP-FLAG}su(Hw)attP1/ P{GMR64C04-GAL4}attP2<br>w[1118] P{y[+t7.7] w[+mC]=hs-FLPG5.PEST}attP3/ w[1118]; +/+; P{y[+t7.7]<br>w[+mC]=10xUAS(FRT.stop)myr::smGdP-OLLAS}attP2 PBac{y[+mDint2]<br>w[+mC]=10xUAS(FRT.stop)myr::smGdP-HA} VK00005 P{10xUAS(FRT.stop)myr::smGdP-V5-THS-10xUAS(FRT.stop)myr::smGdP-FLAG}su(Hw)attP1/ P{GMR21D12-GAL4}attP2 |
| **Figure 3C-F** | P{UAS-RedStinger}3, w[1118]/w[1118]; P{lexA-2xhrgFP.nls}2b/+; Mi{Trojan-<br>lexA:QFAD.0}ChAT[MI04508-TlexA:QFAD.0]/ P{GMR73D10-GAL4}attP2<br>P{UAS-RedStinger}3, w[1118]/ w[1118]; P{lexA-2xhrgFP.nls}2b/+; Mi{Trojan-<br>lexA:QFAD.0}ChAT[MI04508-TlexA:QFAD.0]/ P{GMR64C04-GAL4}attP2<br>P{UAS-RedStinger}3, w[1118]/ w[1118]; P{lexA-2xhrgFP.nls}2b/+; Mi{Trojan-<br>lexA:QFAD.0}ChAT[MI04508-TlexA:QFAD.0]/ P{GMR21D12-GAL4}attP2 |
| **Figure 3G** | w[1118]; P{JFRC7-20XUAS-IVS-mCD8::GFP} attp40/+; P{GMR73D10-GAL4}attP2/+<br>w[1118]; P{JFRC7-20XUAS-IVS-mCD8::GFP} attp40/+; P{GMR64C04-GAL4}attP2/+<br>w[1118]; P{JFRC7-20XUAS-IVS-mCD8::GFP} attp40/+; P{GMR21D12-GAL4}attP2/+ |
| **Figure 4A** | W[1118]/+; P{20xUAS-IVS-GCaMP6f} attP40 /+; P{GMR73D10-GAL4}attP2/P{w[+mc]=UAS-<br>tdTomato} |
| **Figure 4B** | W[1118]/+; P{20xUAS-IVS-GCaMP6f} attP40 /+; P{GMR64C04-GAL4}attP2/P{w[+mc]=UAS-<br>tdTomato} |
| **Figure 4C** | W[1118]/+; P{20xUAS-IVS-GCaMP6f} attP40; P{GMR21D12-GAL4}attP2/P{w[+mc]=UAS-<br>tdTomato} |
| **Figure 5A, D-E.** | W[1118]/+; P{20xUAS-IVS-GCaMP6f} attP40 /+; P{GMR73D10-GAL4}attP2/P{w[+mc]=UAS-tdTomato} |
| **Figure 5B** | W[1118]/+; P{20xUAS-IVS-GCaMP6f} attP40 /+; P{GMR64C04-GAL4}attP2/P{w[+mc]=UAS-tdTomato} |
| **Figure 5C** | W[1118]/+; P{20xUAS-IVS-GCaMP6f} attP40; P{GMR21D12-GAL4}attP2/P{w[+mc]=UAS-tdTomato} |
| **Figure 6B, E-G** | W[1118]/+; P{20xUAS-IVS-GCaMP6f} attP40 /+; P{GMR64C04-GAL4}attP2/P{w[+mc]=UAS-tdTomato} |
| **Figure 6D** | W[1118]/+; P{20xUAS-IVS-GCaMP6f} attP40 /+; P{GMR64C04-GAL4}attP2/P{w[+mc]=UAS-tdTomato}<br>W[1118]/+; P{20xUAS-IVS-GCaMP6f} attP40 /+; P{GMR73D10-GAL4}attP2/P{w[+mc]=UAS-tdTomato}<br>W[1118]/+; P{20xUAS-IVS-GCaMP6f} attP40 /+; P{GMR21D12-GAL4}attP2/P{w[+mc]=UAS-tdTomato} |

